# Unraveling the Genomic and Phylogenetic Complexity of the understudied microfungus *Basidiobolus:* Insights from 19 Newly Sequenced Genomes

**DOI:** 10.1101/2025.03.31.646025

**Authors:** Jasper P. Carleton, Alexander J. Bradshaw, Liam Cleary, Madison R. Hincher, Kathryn Bushley, Javier F. Tabima

## Abstract

*Basidiobolus* is a globally distributed genus of early-diverging fungi within Zoopagomycota, known for its presence in diverse ecological niches ranging from soil and decaying organic matter to vertebrate gastrointestinal tracts. Despite its ecological and medical relevance, the taxonomy and evolutionary relationships within the genus remain poorly resolved due to limited genomic resources. In this study, we present nineteen newly sequenced *Basidiobolus* genomes, expanding the available genomic data. Using short-read Illumina sequencing, assembly, and annotation pipelines, we characterize genic content, assess completeness, and explore biosynthetic gene content across isolates. Phylogenomic analysis reveals two major clades corresponding to *B. meristosporus* and *B. ranarum*, while *B. heterosporus* forms a distinct lineage. Several isolate clusters exhibit deep divergence suggestive of cryptic species, underscoring the need for expanded sampling and taxonomic revision. Functional annotations reveal a rich repertoire of biosynthetic gene clusters, including non-ribosomal peptide synthetases, polyketide synthases, and hybrid clusters, pointing to an underexplored reservoir of secondary metabolite diversity. These findings position *Basidiobolus* as a compelling model for investigating fungal evolution, ecological adaptation, and natural product biosynthesis.

## Introduction

*Basidiobolus* Eidam, a genus of enigmatic fungi within the Zoopagomycota, exhibits a global distribution, thriving in diverse ecological niches. Species of Basidiobolus have been found in organic detritus, including leaf litter and soil (Drechsler, 1964), in the gastrointestinal tract of vertebrates (Gugnani & Okafor, 1980), and are opportunistic pathogens of vertebrates including humans (Gugnani, 1999; Sitterlé et al., 2017; Vanbreuseghem et al., 1978). Despite the genus’s broad presence and ecological versatility, its taxonomy remains poorly resolved. Although fifteen species have been described, only six species have available type cultures. Taxonomic ambiguity within the genus arises from limited knowledge of *Basidiobolus* diversity and is further exacerbated by a scarcity of genomic resources.

*Basidiobolus* genomes hold immense potential for understanding the evolution of chemical diversity in early-diverging fungi. Their genomic architecture includes an array of secondary metabolite (SM) genes, many of which are associated with biosynthetic gene clusters (BGCs) (Tabima et al., 2020, Vargas-Gastelúm et al. 2024). These biosynthetic pathways are hypothesized to contribute to antibiosis, host modulation, and metal sequestration, possibly facilitated by horizontal gene transfer (HGT) from bacteria that coexist in vertebrate gastrointestinal environments (Tabima et al., 2020). Despite the potential for novel secondary metabolites, genomic studies on Basidiobolus remain limited. To date, only three publicly available genomes exist: two identified as *Basidiobolus meristosporus* Drechsler (Chibucos et al., 2016; Mondo et al., 2017), and one as *Basidiobolus heterosporus Drechsler* (Chibucos et al., 2016). Given the high diversity of the genus and its unique biosynthetic potential, the availability of additional genomic resources is critical for exploring its untapped metabolic repertoire.

This study significantly expands genomic representation within Basidiobolus by sequencing, assembling, and annotating nineteen novel isolates. The newly sequenced genomes increase the number of publicly available Basidiobolus genomes from three to twenty-two. Short-read sequencing using Illumina technology was employed to generate high-quality genome assemblies, providing a foundation for comparative genomic analyses. The increased taxon sampling will enable researchers to examine the evolutionary history of Basidiobolus, resolve its taxonomic uncertainties, and investigate the functional roles of its biosynthetic gene clusters.

By addressing the gap in genomic data, this work facilitates a deeper understanding of Basidiobolus as a model system for fungal evolution and secondary metabolism. The expanded genomic dataset lays the groundwork for future studies into host interactions, ecological adaptation, and the potential biotechnological applications of Basidiobolus-derived secondary metabolites. Furthermore, resolving the genetic basis of its unique ecological strategies may provide insights into fungal evolution at the intersection of symbiosis and pathogenicity. Ultimately, the genomic characterization of this enigmatic genus offers a valuable resource for mycologists, evolutionary biologists, and natural product researchers alike.

## Materials and Methods

### Data collection and DNA extraction

Fifteen *Basidiobolus* cultures were acquired from the United States Department of Agriculture’s Agricultural Research Service’s Entomopathogenic Fungi (ARSEF) culture collection in Ithaca, NY (Table 1). Additionally, four specimens were isolated from fecal pellets obtained from amphibians in Worcester County, Massachusetts, using a traditional canopy method for single spore isolation as described by Hincher et al. (2025).

**Table 1.** Strain information and summary metrics for sequence, assembly and quality control of *Basidiobolus* genomes.

| Strain ID | Strain Host | Strain locality | Strain year of Isolation | Raw read count | Aproximate Coverage compared to reference | N50 | Scaffold number | Total Length | Average Contig | Number of gaps | Total gap length | BUSCO (C%) | BUSCO (D%) | BUSCO (F%) |
| --- | --- | --- | --- | --- | --- | --- | --- | --- | --- | --- | --- | --- | --- | --- |
| ARSEF_270 | *Bufo marinus* | USA | 1978 | 39052410 | 65 | 16,234 | 13,781 | 57,360,800 | 4,162 | 862 | 75,115 | 74.8 | 19.8 | 15.2 |
| ARSEF_9887 | *Dyscophus guineti* | Switzerland | 2009 | 39016458 | 65 | 5,865 | 23,970 | 64,439,659 | 2,688 | 817 | 69,046 | 62.3 | 17 | 19.9 |
| ARSEF_9890 | *Hylarana erythraea* | Malaysia | 2009 | 37618954 | 63 | 17,561 | 15,644 | 69,414,244 | 4,437 | 635 | 62,719 | 82.7 | 23.5 | 12.5 |
| ARSEF_9891 | *Hylarana labialis* | Malaysia | 2009 | 35525306 | 59 | 7,134 | 23,511 | 68,729,579 | 2,923 | 693 | 63,699 | 65.5 | 18.4 | 21.8 |
| ARSEF_9892 | *Hylarana glandulosa* | Malaysia | 2009 | 39283862 | 65 | 5,950 | 32,773 | 86,819,798 | 2,649 | 430 | 39,807 | 56.4 | 18.4 | 22.7 |
| ARSEF_9894 | *Bufo bufo* | England | 2009 | 38840514 | 65 | 5,924 | 29,709 | 81,118,883 | 2,730 | 923 | 65,796 | 62.8 | 22.9 | 21.8 |
| ARSEF_9898 | *Bufo bufo* | England | 2009 | 40738096 | 68 | 8,169 | 23,536 | 79,165,723 | 3,364 | 1,075 | 73,477 | 68.6 | 24.1 | 18.1 |
| ARSEF_9899 | *Bufo bufo* | England | 2009 | 39490530 | 66 | 9,361 | 24,963 | 82,929,423 | 3,322 | 441 | 39,995 | 64 | 23.7 | 19.9 |
| ARSEF_9900 | *Discoglossus sp* | Italy | 2009 | 39449118 | 66 | 5,556 | 31,847 | 83,186,951 | 2,612 | 1,213 | 92,745 | 63.1 | 20.8 | 20.4 |
| ARSEF_9901 | *Discoglossus sp* | Italy | 2009 | 44293542 | 74 | 8,419 | 27,588 | 91,543,410 | 3,318 | 798 | 70,594 | 67.8 | 24.2 | 18.6 |
| ARSEF_9903 | *Alytes sp.* | England | 2009 | 43164052 | 72 | 6,955 | 23,896 | 69,009,201 | 2,888 | 740 | 68,873 | 64.5 | 17.6 | 21.2 |
| ARSEF_9904 | *Alytes sp.* | England | 2009 | 38905108 | 65 | 3,206 | 39,129 | 79,801,576 | 2,039 | 1,691 | 105,093 | 59 | 18.5 | 21.7 |
| ARSEF_9908 | *Bufo bufo* | England | 2009 | 49112314 | 82 | 4,792 | 34,177 | 84,632,324 | 2,476 | 214 | 16,358 | 55.3 | 18.5 | 22 |
| ARSEF_9909 | *Tadpole, Alytes sp.* | England | 2009 | 43856332 | 73 | 5,324 | 34,072 | 85,103,423 | 2,498 | 170 | 14,239 | 53.8 | 18.8 | 23.6 |
| ARSEF_9912 | *Alytes sp.* | England | 2009 | 36748884 | 61 | 10,955 | 19,491 | 73,581,624 | 3,775 | 1,166 | 18,001 | 81.3 | 28.2 | 12.8 |
| B_8_4 | *Lithobates catesbeianus* | USA | 2022 | 33783458 | 56 | 11,818 | 18,145 | 63,603,512 | 3,505 | 1,052 | 96,703 | 67.3 | 20 | 19 |
| MH_6_3 | *Lithobates clamitans* | USA | 2022 |  | 64 | 14,908 | 20,979 | 85,826,294 | 4,091 | 964 | 81,679 | 83.5 | 31.4 | 11.7 |
| MH_8_1 | *Lithobates clamitans* | USA | 2022 | 38235252 | 62 | 5,211 | 34,668 | 85,170,883 | 2,457 | 641 | 56,183 | 55.9 | 18.4 | 23.8 |
| MH_9_1 | *Lithobates clamitans* | USA | 2022 | 37303728 | 62 | 5,611 | 31,436 | 79,019,151 | 2,514 | 999 | 92,486 | 62.4 | 22 | 23.4 |

DNA extractions for all nineteen samples were conducted obtained growing the isolates on potato dextrose broth for two weeks, filtered using a vacuum funnel and paper filter, and flash-frozen using liquid nitrogen. The flash frozen tissue was subjected to a CTAB chemical lysis DNA extraction (Carter-House et al., 2020) with the following modifications: the addition of 4uL of RNAse A during the cell lysis step, centrifugation of samples at 14000xg at the washing step, followed by the transfer of the supernatant into new tubes, and the use of .7x volume of isopropanol rather than 1x volume. The purity of the DNA was assessed using the Nanodrop OneC UV-Vis Spectrometer, while dsDNA quantity was assessed using a Qubit 4 Fluorometer. DNA was sequenced on a Novogene corp. Illumina NovaSeq 2×150 S4 flowcell. Raw sequencing results are reported in Table 1.

### Internal transcribed spacer amplification and sequencing

gDNA from each isolate was used for polymerase chain reaction (PCR) of the fungal internal transcribed spacer (ITS) universal barcode region (Schoch *et al*., 2012). Successful amplification was confirmed using a 1% gel electrophoresis and PCR products were cleaned using ExosSAP-IT (Applied Biosystems/ThermoFisher Scientific, Massachusetts, USA). The ITS PCR products were sent for Sanger sequencing at Psomagen corporation (Brooklyn, New York). Forward and reverse sequences were combined into a single consensus sequence for each sample using sangeranalyseR version 1.10.0 (Chao et al., 2021) for all samples and deposited in NCBI GenBank (Table 1).

### Genome assembly and functional annotation

Raw read sequences were first trimmed using trimmomatic (version)(Bolger *et al*., 2014), followed by genome assembly performed using the Automatic Assembly for the Fungi (AAFTF v. 0.4.1) package (Stajich & Palmer, 2023) pipeline. To assess the completeness of the genome, BUSCO v5.83 was performed using the fungi_odb12 dataset (Simão *et al*., 2015) and MetaEuk (Levy Karin *et al*., 2020) prediction method due to lack of *Zoopagomycota* training datasets available for Augustus. Annotations of the genomes were performed using Funannotate V1.8.17 (Palmer & Stajich, 2020)as follows: Assemblies were first cleaned by removing contigs shorter than 500bp, with repeats masked using RepeatMasker V4.0 (Smit et al. 2015). Protein predictions were then performed using four separate prediction methods (Augustus, GlimmerHMM, SNAP, GeneMark-ES)(Korf, 2004; Majoros *et al*., 2004; Stanke *et al*., 2006; Borodovsky & Lomsadze, 2011) using the gene models and predicted protein sequences from the *B. mersitosporus* reference genome (Mondo et al. 2017) as initial evidence. High quality predictions where then determined using EvidenceModeler (Haas *et al*., 2008) with equal weight provided to all prediction methods. Functional annotation was performed using InterProScan5 v 5.59_91.0 (Jones *et al*., 2014) and eggnog-mapper v 2.1.12 (Cantalapiedra *et al*., 2021). BGC annotation was performed with antiSMASH v 7.1.0 (Blin *et al*., 2023). Finally, secreted and transmembrane protein prediction was performed with Phobius v1.01 (Käll *et al*., 2004) and the fast algorithm of SignalP6 (Teufel *et al*., 2022).

### Phylogenomic reconstruction

To produce a phylogenomic data set, we chose to use complete BUSCO genes (fungal_odb12 database) from our samples and the publicly available genomes of *B. meristosporus* CBS 931.73 (Mondo et al. 2018), *B. heterosporus* (Chibucos et al., 2016) and the genome of *B. ranarum* AG-B5 available in NCBI (GCA_040143965.1). Multiple sequence alignment (MSA) of each BUSCO gene was performed using MAFFT v7.525 with the flags --maxiterate 1000 – localpair --reorder (Katoh, 2002). MSA were filtered using the “align filter” function of SEGUL 0.22.1 (Handika & Esselstyn, 2022). Filtering of BUSCO genes was performed using multiple minimal taxon percentage (−-npercent) to attain matrix completion of each gene using two conservative percentage thresholds of 85% and 95%. Pre and post-filtering stats for the MSAs were performed using the “align summary” function of SEGUL (supplementary data). Concatenated and partitions files were then generated using SEGUL “align concat” for downstream branch length estimation. Individual gene trees were created for both filtering schemes using IQTREE v2.4.0 (Minh *et al*., 2020) using ModelFinder (−m MFP)(Kalyaanamoorthy *et al*., 2017) and 1,000 ultrafast bootstrap (−bb 1000) replicates optimized using the --bnni flag (Hoang *et al*., 2018). Genes trees were concatenated and used for summary coalescent analysis using Accurate Species Tree EstimatoR (ASTER⍰) v1.19 (https://github.com/chaoszhang/ASTER), utilizing the ASTRAL-IV model (Zhang & Mirarab, 2022) due to the HGT activity theorized to be occurring in *Basidiobolus*. Branch lengths for the summary coalescent species tree were then estimated using IQ-TREE and with the concatenated and partitions file generated for each filtering scheme. Both filtering schemes generated final trees with identical topology (supplementary data). Tree visualization was done using FigTree 1.4.4 (http://tree.bio.ed.ac.uk/software/figtree/). The tree was rooted on the midpoint. Colors and manual editing of the tree was performed in Adobe Illustrator.

## Results

### Genome Assembly and Completeness

The genome assemblies of the nineteen newly sequenced Basidiobolus strains exhibit a broad range in size, spanning from 51.8 Mbp (ARSEF 270) to 96.1 Mbp (ARSEF 9901) (Table 1). These genome sizes align with previously published reference genomes, including the hybrid assembly *of B. meristosporus* at 89.49 Mbp (Mondo et al., 2017), the short-read assembly of *B. heterosporus* at 44.1 Mbp (Chibucos et al., 2016), and the *B. ranarum* AG-B5 sample from NCBI at 65 Mbp.

Completeness of the genome assemblies, as assessed using BUSCO, ranged from 54% (ARSEF 9909) to 82% (MH 6.3), which is a range comparable to the published and available genomes, with 90% completeness for *B. meristosporus*, 80% for *B. ranarum AG-B5*, and 56.0% for *B. heterosporus*. These discrepancies highlight the necessity for more comprehensive genomic databases to accurately assess genome completeness in early-diverging fungal lineages.

### Gene Annotations and Functional Content

The number of predicted genes per genome ranged from 14,888 (ARSEF 270) to 24,464 (ARSEF 9909) (Table 2), showing variability like other *Basidiobolus* species. These counts are in line with prior genomic studies, which reported 16,111 genes for *B. meristosporus*, 9,331 genes for B. heterosporus (Mondo et al., 2017; Chibucos et al., 2016) and the predicted 18,635 genes for *B. ranarum AG-B5*. Functional annotation using Funannotate revealed genes linked to diverse metabolic and cellular functions, including carbohydrate-active enzymes (dbCAN), general protein families (eggNOG), proteases (MEROPS), and predicted secretome and transmembrane proteins similar to the previously reported *Basidiobolus* genomes (Mondo et al., 2017; Chibucos et al., 2016).

**Table 2.**
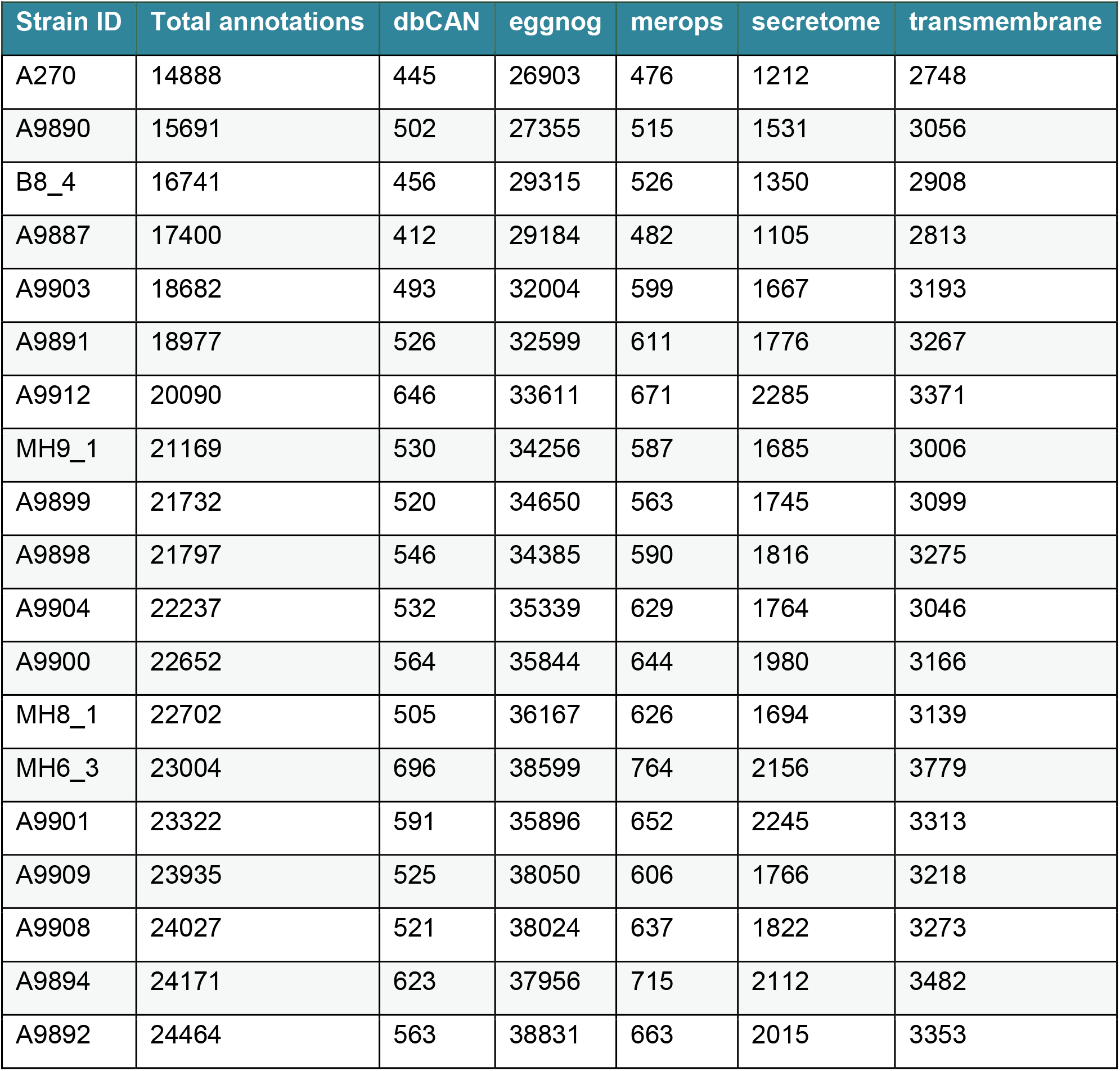
Summary metrics for functional annotation of assembled *Basidiobolus* genomes.

### Biosynthetic Gene Clusters and Secondary Metabolism

The antiSMASH pipeline identified a notable diversity of biosynthetic gene clusters (BGCs), ranging from 30 (ARSEF 270) to 118 (ARSEF 9912) (Table 3). These BGC counts significantly exceed those previously reported for *B. meristosporus* (44 BGCs) and *B. heterosporus* (23 BGCs), reinforcing the hypothesis that *Basidiobolus* species are rich reservoirs of secondary metabolites.

**Table 3.** Summary metrics for secondary metabolism genes from assembled *Basidiobolus* genomes.

| Strain ID | NRPS | NRPS-like | T1PKS | Hybrid | Terpene | Others | Total |
| --- | --- | --- | --- | --- | --- | --- | --- |
| A270 | 17 | 4 | 0 | 0 | 6 | 3 | 30 |
| A9887 | 13 | 8 | 1 | 0 | 6 | 3 | 31 |
| A9890 | 36 | 9 | 3 | 2 | 8 | 3 | 61 |
| A9891 | 49 | 14 | 4 | 0 | 6 | 2 | 75 |
| A9892 | 76 | 25 | 1 | 1 | 6 | 2 | 111 |
| A9894 | 46 | 19 | 2 | 1 | 5 | 2 | 75 |
| A9898 | 30 | 9 | 0 | 0 | 4 | 3 | 46 |
| A9899 | 28 | 9 | 0 | 0 | 6 | 2 | 45 |
| A9900 | 72 | 24 | 2 | 0 | 7 | 2 | 107 |
| A9901 | 75 | 20 | 3 | 0 | 7 | 2 | 107 |
| A9903 | 43 | 11 | 2 | 0 | 6 | 4 | 66 |
| A9904 | 67 | 24 | 2 | 0 | 5 | 2 | 100 |
| A9908 | 50 | 12 | 1 | 0 | 6 | 2 | 71 |
| AA909 | 56 | 9 | 1 | 0 | 7 | 2 | 75 |
| A9912 | 85 | 21 | 3 | 1 | 6 | 4 | 120 |
| B8.4 | 25 | 7 | 0 | 1 | 8 | 2 | 43 |
| MH6.3 | 67 | 18 | 1 | 0 | 9 | 6 | 101 |
| MH8.1 | 57 | 16 | 0 | 1 | 6 | 2 | 82 |
| MH9.1 | 50 | 11 | 0 | 0 | 5 | 3 | 69 |

Among the biosynthetic clusters, non-ribosomal peptide synthetases (NRPS) were the most abundant, with up to 76 clusters in ARSEF 9892. Hybrid clusters, including NRPS-like and polyketide synthase (PKS) pathways, were also identified, suggesting potential for novel bioactive compound discovery. The presence of numerous terpene and other secondary metabolite clusters further underscores the biosynthetic versatility of these fungi.

Collectively, these findings highlight *Basidiobolus* as a promising model for studying fungal secondary metabolism, with significant potential for biotechnological and pharmaceutical applications.

### Phylogenomic reconstruction

Phylogenomic reconstruction was performed using complete BUSCO gene datasets derived from the conserved core of fungal genes (fungi_odb12) dataset. Due to the overall variability of BUSCO completeness, we chose to be conservative with our filtering by using an 85% (p85) and 95%(p95) minimal taxon percentage threshold. 121 complete BUSCO genes were recovered from p85, and 86 from p95. This corresponded to input gene matrices with 13.72 %(p85) and 13.71 %(p95) missing data, and 39,067 (p85) and 30,112 (p95) parsimony informative sites (supplementary data). Phylogenomic reconstruction of both datasets produced trees with identical topology, and p85 was chosen to be used for downstream analysis and visualization (Figure 1).

**Figure 1.**
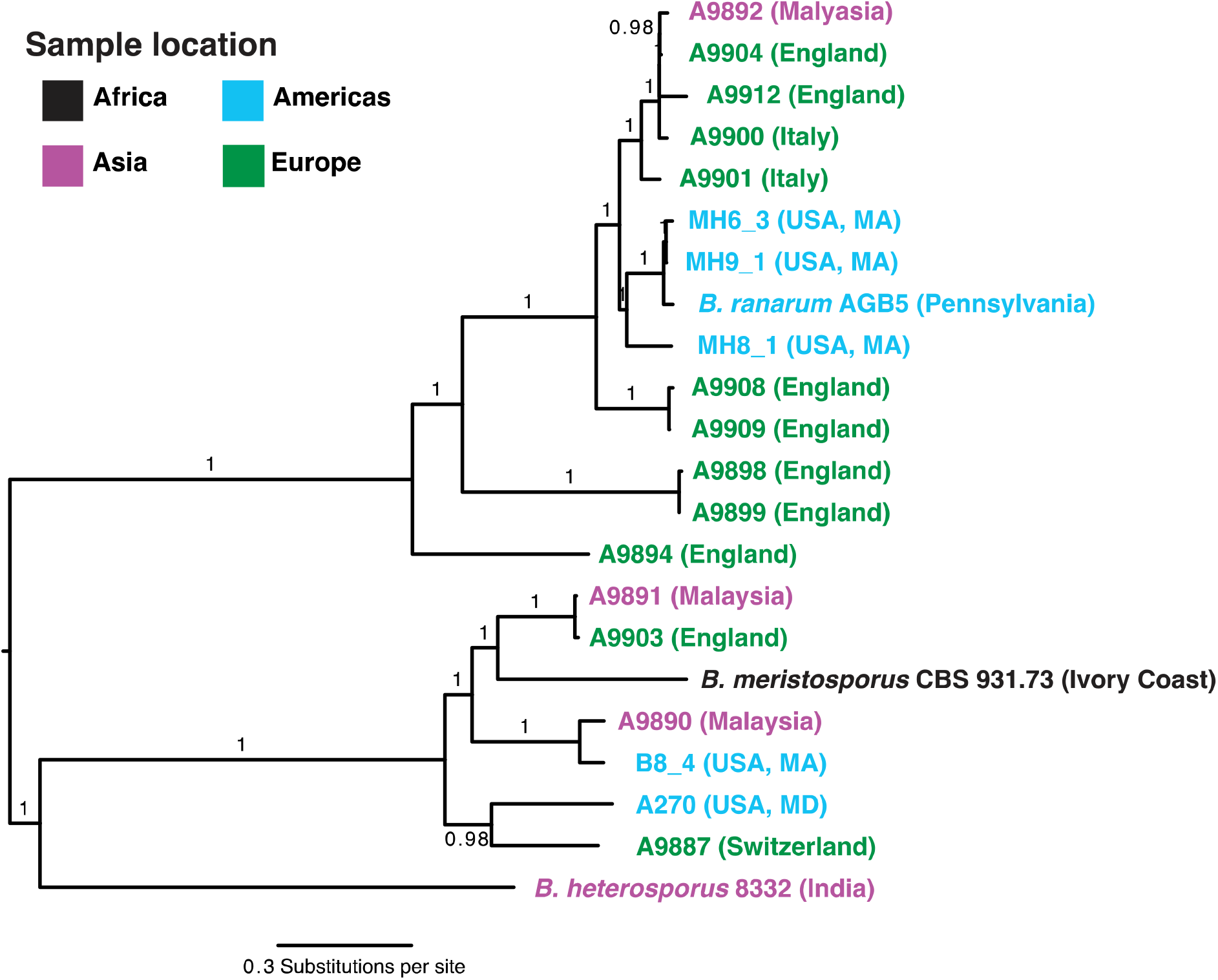
Phylogenomic reconstruction of *Basidiobolus* isolates using complete BUSCO genes, and a minimal taxon percentage threshold of 85%. Tiple labels correspond to isolate designation as well as locality of isolation and are colored according to their continental distribution. Branch labels correspond to bootstrap support values.

Phylogenomic reconstruction revealed two major clades in *Basidiobolus*, one that groups samples to *B. meristosporus* and the other that groups isolates to *B. ranarum AG-B5* (Figure 1).

## Discussion

While genomic sequencing becomes cheaper and more accessible than ever, many organisms, especially cryptic fungi, remain overwhelmingly under-sampled. This underrepresentation hinders our ability to holistically understand fungal diversity and evolution, which already suffers from taxonomic bias in biodiversity studies. *Basidiobolus* presents a unique ecological and evolutionary model due to its presence in diverse environments, ranging from organic detritus to vertebrate gastrointestinal tracts. However, despite its broad distribution, much remains unknown about its evolutionary history, ecological roles, and functional genetics.

Our study significantly expands the genomic dataset available for *Basidiobolus*, providing insights into the genus’s phylogenetic structure, biosynthetic potential, and genetic diversity. The phylogenomic analysis of *Basidiobolus* revealed significant genetic divergence within the genus, highlighting unresolved taxonomic relationships and potential cryptic species. The discovery of distinct clades suggests that *Basidiobolus* harbors greater genetic diversity than previously recognized, reinforcing the need for expanded sampling and molecular analyses. These findings emphasize the limitations of traditional classification methods in capturing the full evolutionary complexity of early-diverging fungi. Additionally, the placement of *B. heterosporus* as phylogenetically distant from both *B. meristosporus* and *B. ranarum* raises questions about its evolutionary trajectory within the genus, warranting further investigation into its genetic and phenotypic distinctiveness.

Beyond taxonomy, the phylogenomic reconstruction also provides insights into the potential dispersal mechanisms of *Basidiobolus*. The close genetic relationship between geographically distant isolates suggests that this genus may possess effective means of long-range dispersal, possibly through animal hosts, environmental transport, or other ecological interactions. This raises important questions about the biogeography and ecological adaptation of *Basidiobolus* species, particularly in relation to their role as both environmental fungi and vertebrate-associated microbes. Future research should integrate genomic, ecological, and experimental approaches to elucidate the mechanisms driving *Basidiobolus* distribution and its evolutionary history. The placement of *B. heterosporus* however, appears to be distant to both these clades. Some cryptic species may be determined in this tree, such as the clade comprising the samples A270 and A9887, the clade that groups A9899 and A9898, and the clade that includes A9908 and A9909. However, much more sampling is needed to determine the taxonomic status and diversity of these clades for a formal species description. Finally, the phylogenetic reconstruction shows that some isolates of *Basidiobolus* that are closely related such as B8_4 and A9890 from very distant geographic regions, indicating that *Basidiobolus* may have some mechanisms of long-range dispersal that need to be further clarified.

A key aspect of *Basidiobolus* biology is its rich secondary metabolite (SM) profile. Our study identified a broad range of biosynthetic gene clusters (BGCs), with some isolates possessing over twice the number of BGCs found in previously sequenced Basidiobolus genomes. The presence of diverse non-ribosomal peptide synthetases (NRPS), polyketide synthases (PKS), and hybrid clusters indicates that *Basidiobolus* may be a valuable source of novel bioactive compounds. Given that secondary metabolites play crucial roles in microbial interactions, host association, and environmental adaptation, future research should focus on characterizing these metabolites to assess their antimicrobial, antifungal, or symbiotic properties. The high prevalence of putative horizontally acquired genes within the BGCs further suggests that *Basidiobolus* may have evolved its metabolic versatility through genetic exchange with bacteria in the vertebrate gut microbiome. Understanding the functional significance of these genes could provide insight into the adaptive strategies that have shaped Basidiobolus evolution.

While our study provides new genomic resources for *Basidiobolus*, it also highlights certain challenges in fungal genome assembly and annotation. The variation in BUSCO completeness scores across isolates suggests that genome quality varies depending on sequencing coverage, assembly methodology, and the limitations of available fungal reference databases. The variation in BUSCO scores could be attributed to differences in sequencing coverage, assembly quality, or limitations in the available *fungi*_*odb12* dataset, which lacks close relatives to *Zoopagomycota* fungi like *Basidiobolus*. The development of more comprehensive phylogenomic databases for early-diverging fungi would enhance the accuracy of future studies. Additionally, transcriptomic and proteomic analyses could complement genome sequencing to improve gene annotations and functional predictions.

Our findings underscore the importance of continued genomic exploration in early-diverging fungi. By expanding the available genomic data, we lay the groundwork for future research into *Basidiobolus* biology, ecological roles, and secondary metabolism. This study contributes to a broader understanding of fungal evolution and supports the use of Basidiobolus as a model system for investigating genetic diversity, biosynthetic potential, and host interactions. Further functional studies are needed to unravel the ecological significance of its diverse metabolic capabilities, and to assess the potential applications of its secondary metabolites in biotechnology and medicine.

## Data Availability Statement

All data for this report can be found on the NCBI BioProject number PRJNA1214880, including raw reads deposited in the NCBI short read archive (SRA), genome assemblies, annotations and ITS sequences, with accession reported in Table 1. All supplementary data including raw trees and alignment statistic files can be found at https://github.com/TabimaLab/Basidiobolus_Carleton_2025

## Notes

https://github.com/TabimaLab/Basidiobolus_Carleton_2025

### Competing Interest Statement

The authors have declared no competing interest.

